# The statistical nature of geometric reasoning

**DOI:** 10.1101/183152

**Authors:** Yuval Hart, Moira R. Dillon, Andrew Marantan, Anna L. Cardenas, Elizabeth Spelke, L. Mahadevan

## Abstract

Geometric reasoning has an inherent dissonance: its abstract axioms and propositions refer to infinitesimal points and infinite straight lines while our perception of the physical world deals with fuzzy dots and curved stripes. How we use these disparate mechanisms to make geometric judgments remains unresolved. Here, we deploy a classically used cognitive geometric task - planar triangle completion - to study the statistics of errors in the location of the missing vertex. Our results show that the mean location has an error proportional to the side of the triangle, the standard deviation is sub-linearly dependent on the side length, and has a negative skewness. These scale-dependent responses directly contradict the conclusions of recent cognitive studies that innate Euclidean rules drive our geometric judgments. To explain our observations, we turn to a perceptual basis for geometric reasoning that balances the competing effects of local smoothness and global orientation of extrapolated trajectories. The resulting mathematical framework captures our observations and further predicts the statistics of the missing angle in a second triangle completion task. To go beyond purely perceptual geometric tasks, we carry out a categorical version of triangle completion that asks about the change in the missing angle after a change in triangle shape. The observed responses show a systematic scale-dependent discrepancy at odds with rule-based Euclidean reasoning, but one that is completely consistent with our framework. All together, our findings point to the use of statistical dynamic models of the noisy perceived physical world, rather than on the abstract rules of Euclid in determining how we reason geometrically.

## Introduction

How do we combine the abstract form of geometry with the physicality of the objects and shapes we see in the world around us? Historically, philosophers from Plato^1^ to Descartes^2^ and Kant^3^, have argued that idealized, abstract geometric entities exist innately in all humans. More recently, scientists like Helmholtz^4^ and Poincaré^5^ have argued that inherently noisy perceptual experience may instead build and shape geometric reasoning. These days, work in cognitive science continues to study the origins and nature of our geometric capacities using philosophical^6^, cross-cultural^7–9^, developmental^10^, and neuro-cognitive approaches^11^.

Previous studies suggested a dominant role for Euclidean rules as the source of our geometric intuitions. For example, when Amazonian adults with no formal education in geometry^8^ were asked to locate a missing vertex in a triangle completion problem, they produced responses that were similar to those of formally educated adults and roughly captured Euclid’s 32^nd^ proposition, that the internal angles of a triangle sum to a constant value, regardless of triangle size. This finding suggested that abstract geometric reasoning is present in all humans, regardless of formal education^7–9,12^, in accord with the thesis that untutored human geometric reasoning is Euclidean, abstract, and perhaps rule-based, i.e. it overrides the physical context in which geometric problems are presented and solved.

However, even educated adults who have mastered the high school curriculum in geometry, do not invoke simple geometrical rules related to changes of square size and scale^13^. To reconcile these differences requires a systematic exploration of the errors made, but this was not possible in previous studies because the triangular stimuli used varied only two-fold in overall size, a variation in the concrete nature of the completion task that is not sufficient to reveal how participants engage with both visual perception and abstract reasoning.

Here, we explore the statistics of errors^14–20^ associated with geometric judgements to infer the underlying perceptual mechanisms. With that aim, we conducted a large-scale triangle completion task, asking participants to locate the missing vertex over a 75-fold range of triangle sizes given the positions and sizes of the other two corners (see Fig 1A). The statistics of participants’ response naturally lead to a quantitative model that balances local and global cognitive demands. Our model makes testable predictions on two related geometric reasoning triangle completion tasks – a perceptual task where participants had to quantify visually the size of the missing third angle, and a categorical task, that asks for qualitative answers about triangle shape (angle) changes.

**Figure 1:**
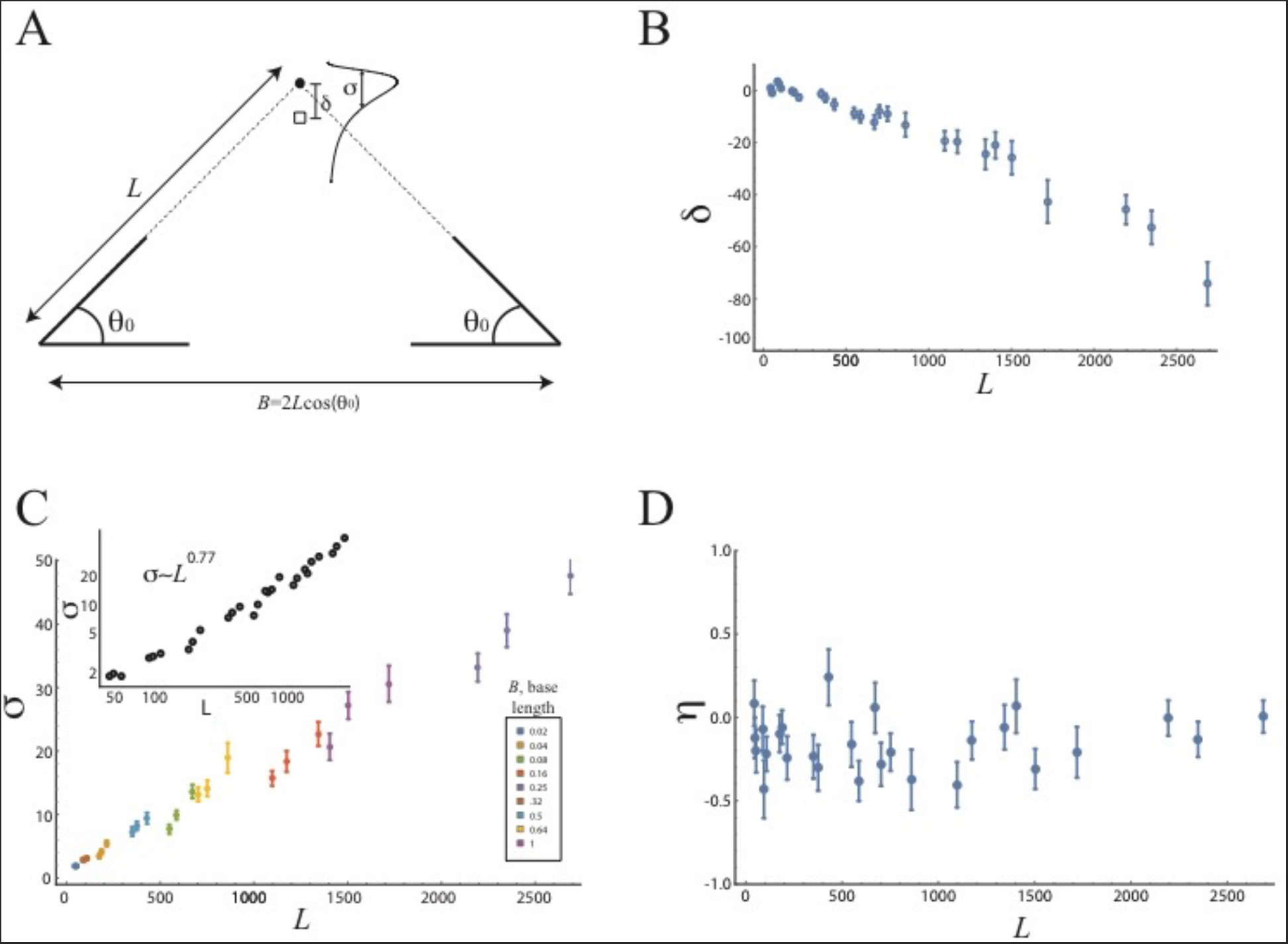
Statistics of the localization of a missing vertex in triangle completion experiment do not follow Euclidean geometry rules. **A)** We measured the distribution of participants’ responses to the location of a triangle’s missing vertex over exemplars that varied by ~75 fold in triangle side-length. Base length values ranged from 0.02 to 1 (where 1 equals 1900 pixels — see Methods) and angles values were 30°, 36°, and 45°. **B)** The mean deviation (*δ*) from the true y-coordinate location of the missing vertex to the mean of participants’ responses as a function of triangle side-length. This deviation scales linearly with triangle’s side length and is biased toward the base. **C)** The standard deviation of participants’ distribution of responses (σ) as a function of side length. Inset (log transformed): this standard deviation scales sub-linearly with the side length (*σ*~*L*^0.77^, Min-Max values: 0.65-1.04, median exponent: 0.77, 95% CI=[0.73,0.82]) **D)** The skewness of participants’ distribution of responses (*η*) as a function of side length, which is negative, *η*=-0.15 (95% CI=[-0.22,-0.09]), indicating that the distribution of participants’ y-coordinate estimates has a long tail with more responses which are biased toward the base of the triangle.

## Results

### Participants responses in a triangle completion localization task are scale dependent

To understand the link between noisy physical representations and geometric judgments, we asked adult participants (N=40) to position the missing vertex of 15 different fragmented triangles (each presented 10 times, all with their base on the x-axis) projected on a large screen. This task allows us to analyze the effects of triangle size on the mean, variance, and skewness of the distribution of responses (Fig 1A). We note that a use of Euclidean rules in this task would generate a linear scaling for the standard deviation with triangle side-length.

Contrary to these expectations, we found that the y-coordinate location estimates were biased toward the base of the triangle, and that this bias increased linearly with triangle side length (Fig. 1B). Significantly, the standard deviation of the y-coordinate location estimates scaled sub-linearly with side length, *σ*~*L*^0.77^ (median exponent: 0.77, 95% CI=[0.73,0.82], Fig 1C and Fig S1), and the response distribution had a negative skewness (*η*=-0.15, 95% CI=[-0.22,-0.09]; Fig 1D). The x-coordinate location estimates also showed a sub-linear scaling of its standard deviation, with no systematic directional bias through most of the triangle size range, although the errors were 4-fold lower in magnitude (SI, Fig S2).

To confirm these initial findings, we repeated the task with a group of online participants (N=100, SI, Fig S3) and found a similar y-coordinate bias toward the base of the triangle and a sub-linear scaling of its standard deviation, *σ*~*L*^0.65^ (SI, Fig S3). The x-coordinate showed no systematic bias in triangle size and sub-linear scaling of its standard deviation. A rotated version of the experiment (N=50), with the triangle base at the y-axis, showed similar results to the versions with the base at the x-axis (SI, Fig S4). These results show that the localization and evaluation of corners on triangle completion tasks are not based on Euclidean rules but rather on a dynamic cognitive-perceptual process dependent on the visual perception of the triangle.

### Response statistics suggest a cognitive-perceptual mechanism that balances between local smoothness of angles estimation and their adherence to the base angle

To explain our observations theoretically, we model the cognitive-perceptual process of determining the missing vertex location. We start with a dynamical model for the triangle completion task^21,22^, where people’s extrapolation process is described by lines emanating from each vertex with their local orientation (denoted *θ*) dynamically evolving. The estimated angles of the extrapolated lines evolve in space and time in the presence of noisy estimates until they intersect (Fig 2A). An error-correction of the estimated angle to the base angle is done at time intervals with a time scale marked by *ξ*. Thus, the parameter *ξ* controls much of the statistics of missing vertex estimation. The dynamic equations are (see Fig 2A):

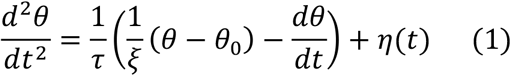

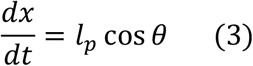

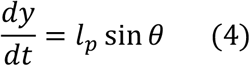

where the parameters of the model are: τ, an inertial relaxation time scale, *l_p_*, a characteristic speed, *ξ*, a time scale for global error-correction, and *η*(*t*) is a noise term with noise amplitude *D*, (⟨*η*(*t*)*η*(*t*′)⟩ = *D* δ(*t* – *t*′)). In addition, the model has a threshold for the x-coordinate distance between the two line extrapolated curves, *ϵ*, which once crossed, ends the process (SI, Figs S5-S7).

**Figure 2:**
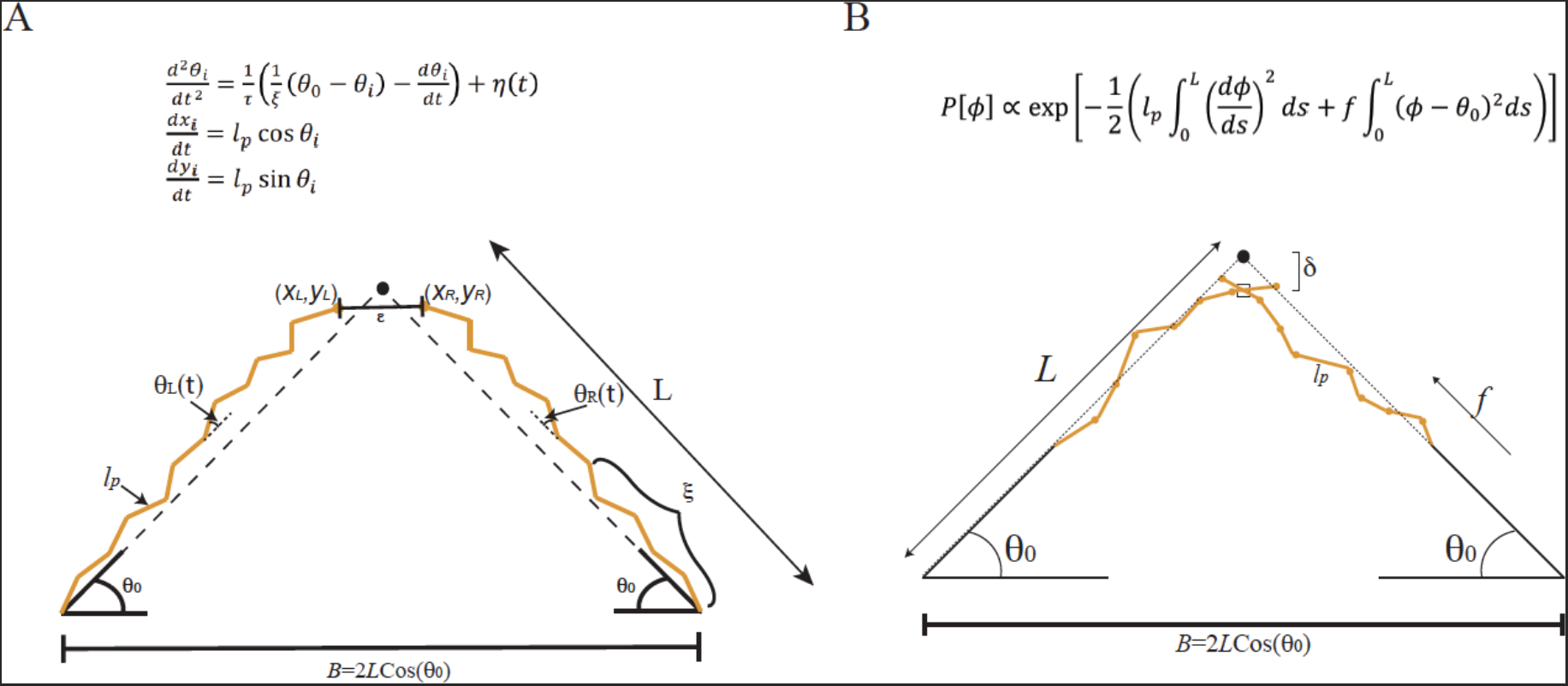
A mathematical model for triangle completion uses correlated random walks that start at the known vertices and progress until they meet. **A)** The schematics of the dynamical model in the triangle completion task. In the dynamic model, the local angle evolves coupled to a noise term as the line trajectory is built. The model parameters are: *l_p_*, a characteristic speed with which the coordinates progress, *ξ*, a time scale for global error-correction (illustrated as number of segments between error-correction events), and η(*t*) is a noise term with noise amplitude *D* (⟨*η*(*t*)*η*(*t*′)⟩ = *D* δ(*t* – *t*′)), not shown in the figure. The stopping criterion threshold is denoted, *ϵ*, and the base angle is denoted by *θ*_0_. The dynamic model converges over many iterations to the statistical model (see SI for further details). **B)** The schematics of the statistical model. Estimate for the location of the missing vertex arises from balancing noisy local orientational order with a global error-correction mechanism. The penalty for local angle deviations relative to the previous segment is characterized by a weight *l_p_*, and the penalty for angle deviations from the global orientation relative to the base angle is characterized by a weight *f*, yielding a correlation length, 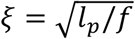, as a fitting parameter.

While this approach provides an appealing picture for the mental process of triangle completion, it depends on 4 different parameters. Next, we describe a statistical approach which has the benefit of just one dominant parameter and which repeated use of the dynamical model converges to.

The statistical approach considers the statistics of a line-like object that is built from small segments that reflect the two competing processes mentioned above – maintaining motion along a straight smooth line and correcting its overall direction given by a base angle value. Together, these two processes yield the following probability for the angle of each segment ϕ(*s*), with *s* characterizing location along the curve, relative to the initial base angle *θ*_0_ (see Fig 2B):

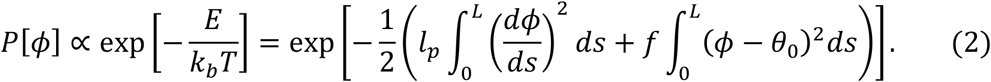

The form of the energy *E* is exactly the Hamiltonian for a model describing semi-flexible polymers^23,24^, and has also been coopted to understand aspects of navigation^25–27^, as it follows from simple invariance properties, balancing the competition between local and global orientational order and straightness. Indeed, the first term reflects the penalty associated with high curvature with a weight known as the persistence length *l_p_*, which defines the magnitude of the local noise in the angle judgment at each segment. The second term reflects the penalty for angle deviations from the initial base angle *θ*_0_, with a weight *f*, which acts as a global error-correction mechanism. The model effectively has one parameter, a correlation length 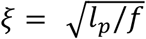, which balances the two competing effects (and which is proportional to the time scale of the dynamical model (1) up to a factor of the speed *l_p_*).

For small *ξ* (relative to the size of the triangle) the error-correction term dominates, and the line extrapolation is a simple random walk with a standard deviation scaling of *σ~L*^0.5^ (see SI). For large *ξ* (relative to the size of the triangle), the local noise term dominates, and line extrapolation is a correlated random walk with a standard deviation *σ~L*^1.5^ (see SI).

The exponent observed in the localization experiment, 0.77, suggests that the error correction mechanism plays a more dominant role in participants’ judgements. Importantly, this signifies better robustness to noise propagation with triangle’s size than the linear dependence produced by the static Euclidean rules. Taking *ξ*~2 times the smallest initial side length (corresponding to an eye movement of ~1 degree) and then varying the side length over almost two decades allows us to capture the mean, standard deviation, and skewness of the distribution of the observed location responses (Fig 3A-B and SI, Figs S8-S11).

**Figure 3:**
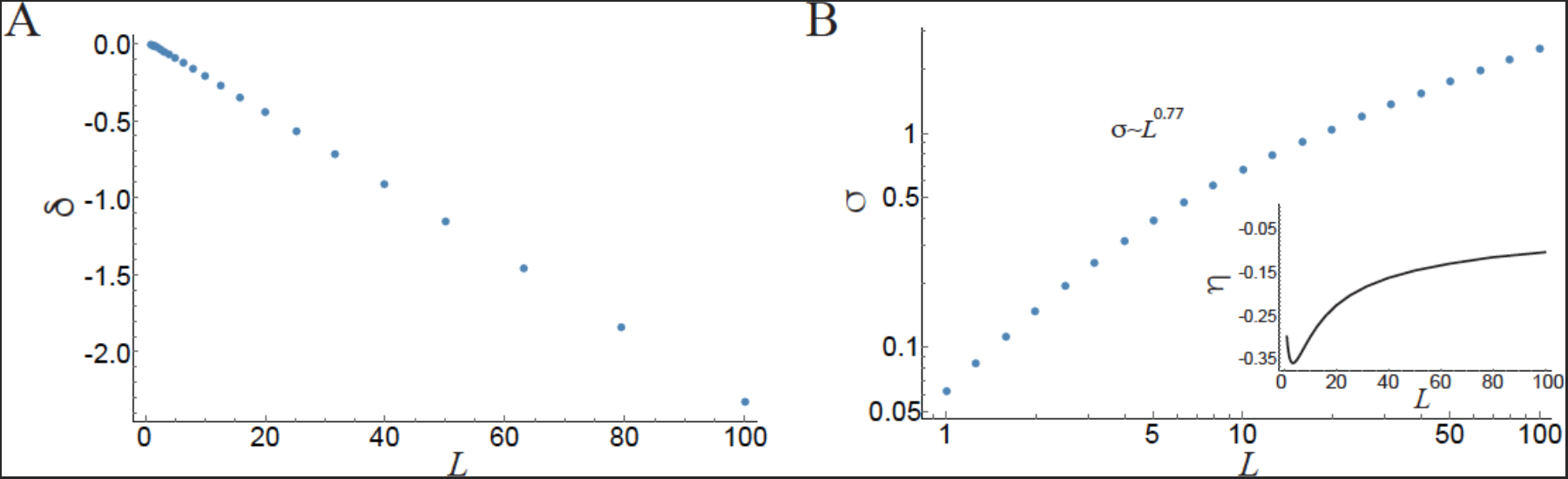
The perceptual-cognitive model captures the behavioral results in the triangle completion localization task. **A)** The results of the statistical model show that y-coordinate location estimates have a bias toward the base of the triangle that increases linearly with the triangle side length. Here, we use *ξ* = 2 times the smallest side length, angle noise level *V*_0_ = (*l_p_f*)^-0.5^ =0.26 (see Methods and SI) and side length varies between 1-100. **B)** Using the same parameters as in A), our results also show the standard deviation of the y-coordinate location estimate varies sub-linearly with side length, *σ*~*L*^0.77^. **Inset**, skewness attains similar negative values. (similar results are obtained by the dynamic model, see SI)

### The model predicts the statistics of angle estimates in a triangle completion task

As a further test of the model, we used it to predict the distribution of participant responses for the magnitude of missing angles, keeping the correlation length fixed. The model predicts that the mean size of the angles would be overestimated and increase as triangle side-length grows, and that the variance of the distribution of angles would decrease as triangle side-length grows (SI, Figs S12-14).

In an online experiment (N=65, SI, Fig S14), a new group of participants were asked to use a goniometer to estimate the missing angle size of 10 instances of 15 different triangles. We found that, as predicted, participants overestimated the missing angle size by an amount that increased with triangle side-length (at large base angles of 45° and 60°, Fig S14) and that the variance of the distribution decreased as triangle side-length grew (Fig S14). These results indicate that our model, derived from location judgments alone, also captured the independently observed responses about the missing angle size. Moreover, the properties of our model indicate that the process for visual triangle completion leads to a systematic inability to preserve the Euclidean properties of planar triangles, i.e., that lines are perfectly straight and that the internal angles sum to a constant. A geometric interpretation of this is that our perception of geometry becomes inherently non-Euclidean owing to the presence of a hidden length scale, the correlation length *ξ*.

### Participants’ responses in a categorical task contradict Euclidean geometry rules

An interesting question that arises is what happens in more abstract reasoning scenarios, that do not necessitate visualization or mental imagery but rather can be solved easily and rapidly by using the powerful Euclidean rules. To test this we conducted a third, categorical reasoning task. In the experiment adult participants on Amazon Mechanical Turk (N=407) were asked in two separate blocks (with no accompanying visual transformations): whether a triangle’s vertex would move up, move down, or stay in the same place after the other two vertices either moved farther apart, closer together, grew in angle size, or shrunk in angle size; and whether the associated angle would get bigger, get smaller, or stay the same size after those same four transformations (totaling 8 multiple choice questions with chance at 33%; Fig 4A). We also measured accuracy and response times of the participants.

**Figure 4:**
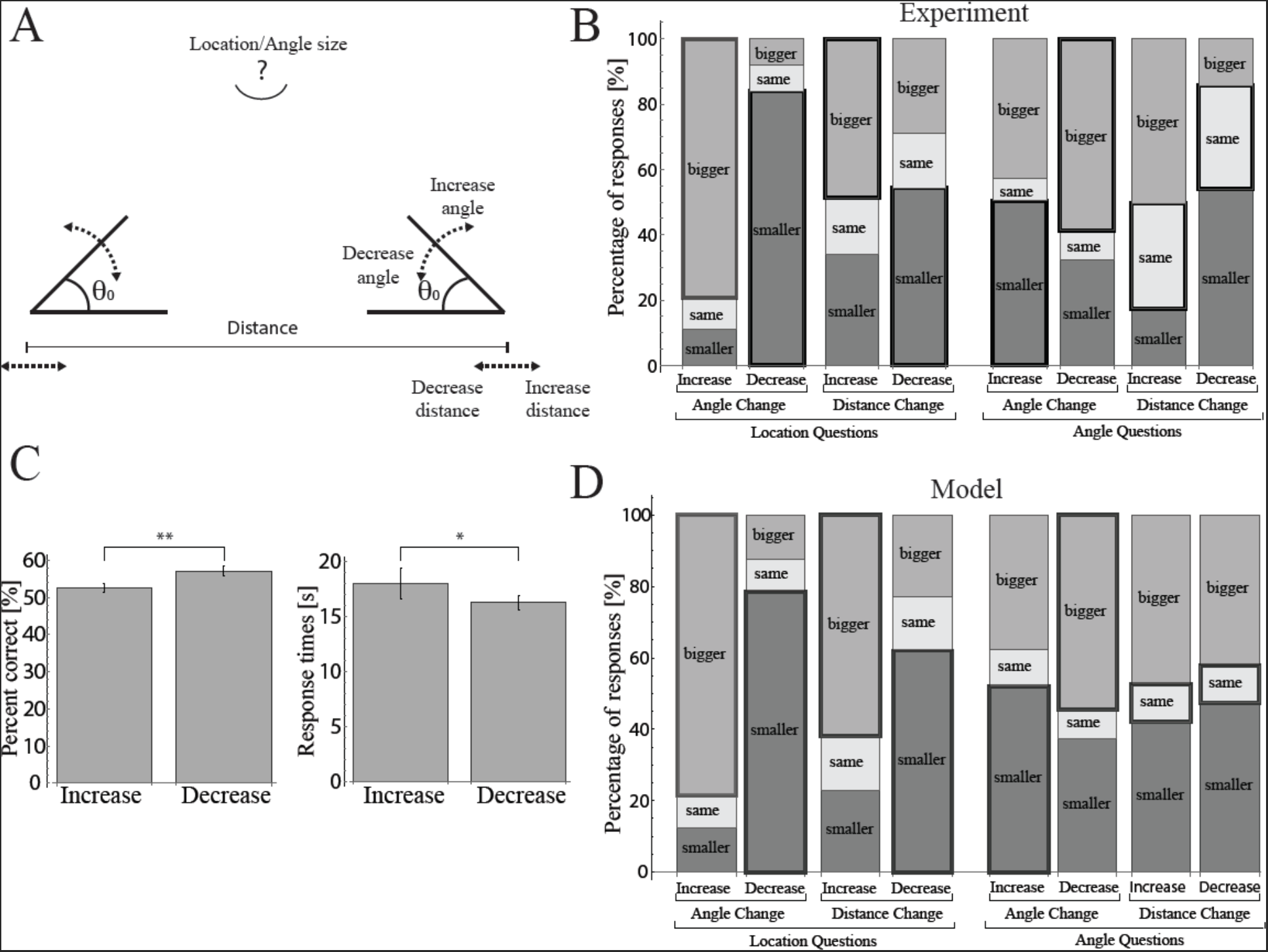
Categorical responses to the geometric reasoning task of triangle completion contradict a use of Euclidean geometry rules. **A)** In separate blocks of questions, participants were asked to judge the change in the location of the missing vertex and the change in the magnitude of the missing angle. **B)** Participants’ responses do not capture Euclidean rules (correct responses are outlined in bold). For example, participants predominantly judge that an angle should scale with the overall size of the triangle (last pair of bars). **C)** Participants’ responses are more accurate and faster when triangles get smaller (the angle sizes or the distance between the vertices decreases) vs. larger (the angle sizes or the distance between the vertices increases). Accuracy: Decrease: Mean±STE=57±1%, Increase: Mean±STE=53±1%, Mann-Whitney test: t(1628^2^)= 1,264,960, p<0.01, effect size =0.05; Response times: Decrease: Mean±STE=16±1 s, Increase: Mean±STE=18±1 s, Mann-Whitney test: t(1628^2^)=1,387,910, p<0.02, effect size = 0.05; *p<0.05, **p<0.01. **D)** The statistical model allows us to explain the predicted responses to the categorical responses associated with the reasoning task shown in Fig. 4B; chi-squared tests with the behavioral data showed values>0.17. A Bayes factor estimate comparing the model with a model of only noise in base angle estimates with similar thresholds showed values of BF > 10^10^ for most questions (with the exception of AID BF=0.08 and ADD BF=3). Model parameters are *ξ*=2, *V*_0_ = 0.4, Th_L_=0.05, Th_A_=0.05, all initial base angles=36 degrees and initial side length distance varied between 2-4 (see Methods and SI).

Participants performed well above chance on location judgments after both angle and distance changes at the other two corners (Fig 4B), as well as on angle judgments after angle changes at the other two corners, but their performance was far from perfect. Most strikingly, however, participants performed at chance levels on angle judgments after changes in the distance between the other two vertices (Fig 4B). Erroneous responses on this question were in direct contradiction with Euclid’s 32^nd^ proposition^28^ that the internal angles of a triangle sum to a constant (e.g., participants responded that the missing angle got bigger when the other two corners maintained their angular measure but moved farther apart; Fig 4B and Fig S15).

To evaluate whether participants relied on rules during this task, we asked whether there were any differences in the responses to the questions referring to transformations that decreased vs. increased the size of the triangle. If participants were using a rule, whereby the size of the third angle is invariant over transformations of scale, then no such differences should be found. However, if participants relied on some perceptual strategy, then we might observe greater success and shorter response times after transformations that decreased the triangle size. Our results show that participants responded more accurately and more quickly after they were asked to make judgments about triangles that got smaller (Accuracy: Mann-Whitney test: t(1628^2^)= 1,264,960, *p*<0.01, effect size =0.05; Response Time: Mann-Whitney test: t(1628^2^)=1,3 87,910, p<0.02, effect size = 0.05; Fig. 4C), consistent with a strategy based on perception or imagery.

### The perceptual-cognitive model captures the behavioral categorical task responses

Although our model relies only on the statistics of the localization task responses, we ask if it can capture the results associated with the categorical geometric reasoning task made by participants. For example, for a question concerning the effect of increasing the distance between the two base vertices on the missing vertex location, we approximated the distribution of the vertex location by a Gamma distribution with the corresponding model values of mean and variance and then compare it to a Gamma distribution with parameters taken from the increased distance condition. Thresholds for the “move up”, “move down” and “same” categories were set according to the ratio of the measured bias and standard deviation of participant location estimates from the lab-based localization task (see Methods and SI). We found that our model produced responses that closely resembled those of our categorical geometric reasoning task for all 8 questions (Chi squared tests, all *ps* > 0.17, Fig 4D vs. Fig 4B). Furthermore, comparing the model results with a model with only noisy estimates of base angles yielded high support for the model (Bayes factors^29^ (BF) for most questions >10^10^, and AID BF=0.08 and ADD BF=3, see Methods and SI). We note that allowing the model’s parameters to vary for each question or adding noise to the base angle estimates improves the model’s fit to results from the categorical geometric reasoning task results (see SI, Figs S16-S19).

## Discussion

Our study provides both behavioral evidence and a theoretical and computational framework that shows how geometric reasoning in adults depends on established cognitive processes that link perception and action: mental simulation of locally correlated motion along a segment of a line^30,31^ and correction of its global direction^21,22^. The former reflects the ability of our visual system to follow a smooth trajectory locally, similar to the well-known gestalt principle of ‘good continuation’^32–36^, and the latter reflects the capacity of short term visual memory to represent global orientation^37–40^. Why should a statistical process be preferred over the use of a simple and static powerful rule? A possible explanation stems from considering the challenges posed by noise in perception. While the static Euclidean rules may provide an accurate and rapid response, under conditions of noisy perception, statistical models can offer a mechanism that is more robust to noise. In lieu of this suggestion is the sublinear dependency of the standard deviation on triangle side-length observed in participants’ estimations. This sublinear dependence is smaller than the expected linear dependency of Euclidean rules which propagate errors in estimation linearly with triangle side-length.

Further work connecting eye tracking and brain activity measurements would serve to understand the determinants of the correlation length ξ that suffices to captures the statistics of our vertex localization task, missing angle estimations and even categorical reasoning.

More generally, our work contributes to accumulating evidence that statistical and dynamical processes are just as critical as static, rule-based thinking in understanding core concepts in psychology, especially in domains like geometry and physics, in which we solve problems by constructing and transforming mental models of the perceived world^41–46^. Since geometry is the bedrock of our physical intuitions, establishing the links between geometrical intuitions and motion assessment and prediction might reveal new insights to the mechanisms governing our perception of the static and dynamic world. For example, crucial skills such as motion assessment and navigation require an accurate geometric perception of space. The predictive strength of a perceptual model for geometry suggests an appealing hypothesis that it is driven by a cognitive advantage associated with the ability to dynamically infer spatial information.

Finally, our work raises the question: how are formalisms of human knowledge discovered and used? Since psychological mechanisms shared by animals, children and adults allow for perception and navigation in uncertain and imperfectly known environments, it is yet to be understood how humans succeeded, over time, to convert reproducible strategies for these tasks into mathematical intuitions and rules.

## Materials and Methods

### Localization of the missing vertex in a triangle completion task - laboratory experiment

In a laboratory experiment, we showed participants 15 different incomplete isosceles triangles 10 times in a random order (for a total of 150 triangles for each participant). Forty participants, divided randomly into two equal-sized groups, were shown triangles of 3 different base angle sizes (30, 36 and 45 degrees) and with 5 different base lengths. Participants in group 1 were shown base lengths of 0.02, 0.08, 0.25, 0.5, and 1. Participants in group 2 were shown base lengths of 0.04, 0.16, 0.32, 0.64, and 1. In both groups, 1 signifies 1900 pixels and is equivalent to 130 cm. Participants sat at a distance of 150 cm from the screen. For each triangle, we asked participants to position a dot in the estimated location of the missing vertex. Before the experiment began, participants had one practice trial, in which the location of the missing vertex was indicated by a dot of a different color, and they were asked to position their dot on the indicated position.

### Localization of the missing vertex in a triangle completion task - online experiment

In an online experiment, we repeated the same task as in the lab experiment with 100 participants, divided into two groups. Base angles were set to 3 different angle sizes 30, 45 and 60 degrees - group 1 (50 participants), and 36, 51 and 66 degrees - group 2 (50 participants), with 5 different base lengths of 0.1, 0.25, 0.5, 0.75 and 1. We used two base length scales for the experiments since triangles would have exceeded the size of the screen with the angle sizes presented in group 2 at the distance scale used in group 1. Group 1 saw a base length scale 900 pixels and group 2 saw a base length scale of 1300 pixels. To match the scales for the two groups we divided the estimates of the second group by a ratio of 13/9.

### Localization of the missing vertex in a triangle completion task - rotated online experiment

We repeated the same task of positioning the missing vertex with a rotated isosceles triangle such that the base of the triangle was on the y axis, on the right side of the screen. 50 participants were shown 3 different base angle sizes (30, 45 and 60 degrees), with 5 different base lengths (0.1, 0.25, 0.5, 0.75 and 1), where a base length of 1 was set to be 1000 pixels.

### Analysis of all behavioral data

All data analyses were done using Mathematica 11.0. The mean deviation from the true location of the missing vertex or the missing angle size, the standard deviation, and the skewness were calculated for each participant and then averaged across participants. Results in the main text show mean±ste.

### Derivation of Y-coordinate distribution mean, variance and skewness

In order to model and predict the quantitative results for the localization task, we assumed the estimated location (*X,Y*) was the average of a right and left triangle’s side trajectory extrapolations (see Fig. S7). We derived analytic expressions for the moments of each side extrapolated trajectory by using 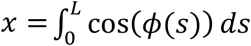 and 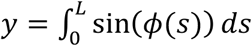 where ϕ(*s*) were taken from the probability distribution: 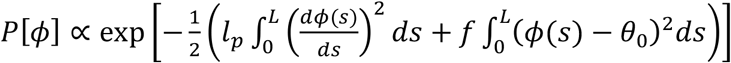. We calculated the bias in the location estimate by subtracting the true location (*y_true_* = *L Sin*[*θ*_0_]) from the mean y-coordinate. The standard deviation was calculated as the square root of the second moment of the distribution of the estimated (*X,Y*) location. The correlation length, 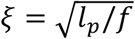, is the dominant parameter setting the scaling exponent between the vertical location standard deviation and the side length. We found a best-fit to the participants’ responses at a value of *ξ* = 2 (where 1 denotes the side-length of the smallest triangle considered), and the side length varies across *L* ∈ [1,100] (a similar range as the experimental setup). A detailed calculation of the moments and sensitivity analysis of model parameters are presented in the SI. And see Fig S20 for the relation between base angle and error in the estimated mean location of the missing vertex.

### Model estimate of the statistics of the missing angle

The magnitude of the missing angle was calculated using estimates of the missing vertex. Given the good fit of the Gamma distribution to participants’ y-coordinate estimates in the localization task (see SI, Fig. S11), we approximated in our model the vertical distribution as a Gamma distribution whose mean and variance were derived from our model’s analytical calculations. The x-coordinate was sampled from a Gaussian distribution with a standard deviation derived from the same analytical calculations. This produced a set of (*X,Y*) locations that was used to derive the estimated missing angle size value. The missing angle size was calculated as: Missing angle size = π - (effective base angle right + effective base angle left). We repeated this process 400 times to produce a distribution of estimated angles per side length and base angle. We then calculated the mean and standard deviation as a function of side length and base angle. We used the same correlation length, *ξ* = 2, and noise levels of *V*_0_ = 0.4.

### Categorical geometric reasoning experiment

In an online experiment (mechanical Turk) we asked participants to answer 8 randomly ordered categorical questions regarding imagined manipulations to triangle size or shape. Participants were presented with the two base vertices of an incomplete isosceles triangle and were asked what would happen to the location (or angle size) of the missing vertex upon an increase (or decrease) of 20% in the distance between (or angle size of) the two bottom vertices. Participants saw the same drawing of a static, fragmented triangle with each question throughout the experiment. In different groups of participants, this accompanying triangle had corners that were either 600 pixels and 240 pixels apart and presented either 36 and 60 degree angles. Each experiment started with a demonstration of what the indicated manipulations to distance and angles of the base corners looked like on a different example triangle. For each imagined manipulation, participants indicated whether the missing corner’s location would move up, move down, or stay in the same place. Similarly, they also indicated whether its angle size would get bigger, get smaller, or stay the same size. Four-hundred-seven participants completed the experiment: 157 females; 427 males; and 3 who did not specify a gender. Participants’ age ranged between 18-72 years, with a median of 31 years. Participants’ years of education ranged between 8-33 years, with a median of 15 years (and see SI, Fig. S1).

### Model estimates of the categorical geometric reasoning task results

The categorical geometric reasoning task of triangle completion challenged participants to compare location or angle size estimates from two triangles, an initial incomplete triangle presented on the screen and an imagined triangle resulting from a specific manipulation (increasing or decreasing the distance between or angle size of the two base angles; see above). We thus compared the model’s predictions for the locations or angles of the initial triangle to the triangle that would result from the indicated manipulation. For example, consider a question about the location change of the missing vertex after an increase of the distance between the two base vertices. We calculated the location estimates of the model for the initial triangle by plugging the model’s predicted mean and variance to a Gamma distribution yielding a sample of 400 estimated locations (see SI, Fig. S11). We then repeated this process for the manipulated triangle. Next, we compared the two samples of location estimates in pairs. For each pair, we calculate the percent change in location, and used a threshold to categorize the answer as “move up”, “move down” or “stays in the same place”: Locations which were 5% higher than the initial estimated location we marked as “move up” 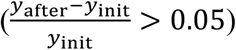. Locations which were 5% lower than the initial estimated location we marked as “move down” 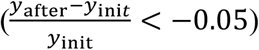. All values in between these two thresholds were marked as “stays in the same place” 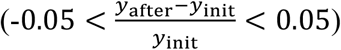. This categorization method was also used with angle questions. The threshold was set by estimating the median coefficient of variation (std/mean) in participants’ answers in the localization task of the missing vertex location estimates (See SI for more details and sensitivity analysis of the thresholding values). For all questions the following parameters were used: correlation length, *ξ* = 1.25, variance of interior angles estimates *V*_0_ = 0,5, initial side-length, L=3.2 for location questions and L=1.25 for angle questions, initial angle=36 degrees, increased angle = 45 degrees, length increase for location questions = 25%, length increase for angle questions = 50%.

### Goodness of fit for the model and the categorical geometric reasoning task results

We used a Chi-Squared test for goodness of fit between the model predictions and the participants’ responses in the categorical geometric reasoning task. These tests did not show a significant difference between the model and participants’ responses. Chi-Squared statistics and p-values were: t(1)=(1.87, 1.87, 1.33, 1.33, 1.33, 0.75, 0.14, 0.14), *p*=(0.17, 0.17, 0.25, 0.25, 0.25, 0.39, 0. 7. 0.7) for VIA,VDA, VID, VDD, AIA, ADA, AID, ADD questions respectively (each condition first letter indicates whether the question concerned vertex location or angle size (V/A), the second letter indicates whether the manipulation concerned increase or decrease in value (I/D) and the third letter indicates whether the manipulation suggested concerned changes to the base angles size or the distance between the base angles (A/D)).

### Bayes factor comparison of the model and a model with only noisy base angle estimates

We used Bayes factor analysis to validate the fit of our model to the categorical geometric reasoning task results. We compared our model with a model that used straight lines with only Gaussian noise in the assessment of the base angle - angles were assumed to be sampled from a Gaussian distribution with the mean set to the base angle and a standard deviation of 5 degrees (see SI for more details). We used the same thresholds for both models (5% change as a detection threshold). The Bayes factor was calculated as

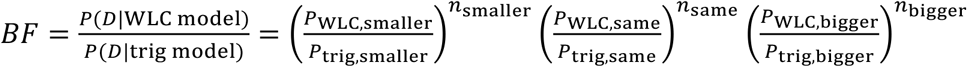

where P_WLC,i_ are the probabilities derived from our model, P_trig,i_ are the probabilities derived from a model with only noisy base angle estimates and ni are the number of such responses in the categorical geometric reasoning task. The BF results were BF=(10^27^, 10^18^, 10^119^, 10^96^, 10^62^, 10^147^, 0.08, 3) for the VIA,VDA, VID, VDD, AIA, ADA, AID, ADD questions respectively, indicating that, for most questions, the WLC model is superior to the simpler, straight-line Euclidean model with noisy base angle estimates.

